# RaMP-DB 2.0: a renovated knowledgebase for deriving biological and chemical insight from genes, proteins, and metabolites

**DOI:** 10.1101/2022.01.19.476987

**Authors:** John Braisted, Andrew Patt, Cole Tindall, Tara Eicher, Timothy Sheils, Jorge Neyra, Kyle Spencer, Ewy A. Mathé

## Abstract

RaMP-DB 2.0 is a web interface, API, relational database and R package designed for straightforward and comprehensive functional interpretation of metabolomic and multi-omic data. Since its first release in 2018, RaMP-DB 2.0 has been upgraded with an expanded breadth and depth of functional and chemical annotation. Content from the source databases (Reactome, HMDB, and Wikipathways) has been updated, and new data types related to metabolite annotations have been incorporated. Structural information incorporated in RaMP-DB 2.0 includes SMILES strings, InChIs, InChIKeys. Chemical classes have been sourced from ClassyFire and LIPID MAPS. Accordingly, the RaMP-DB 2.0 R package has been updated and supports queries on pathways, common reactions, ontologies, chemical classes, and chemical structures. Additionally, RaMP-DB 2.0 now supports enrichment analyses on pathways and chemical classes. Our process for integrating annotations across resources has also been upgraded to lessen the burden of harmonization, thereby supporting more frequent updates. The code used to build all components of RaMP-DB 2.0 is freely available on GitHub at https://github.com/ncats/ramp-db and https://github.com/ncats/RaMP-Backend.

## INTRODUCTION

The impact of metabolomics and multi-omics on biomedical and translational research continues to grow. Multi-omic studies that combine metabolomics data with genomic, transcriptomic, or proteomic data provide additional perspectives that capture the many complex interactions occurring between genes, proteins, and metabolites^1–4^. However, interpretation of multi-omic data raises many hurdles for researchers. Challenges associated with multi-omic integration include the large variety of identifier types for metabolites, genes, and proteins, the scarcity of up-to-date comprehensive and integrated gene/protein and metabolite annotation sources, and the tools to work across these omics types. With these issues in mind, we created RaMP-DB 2.0, the relational database of metabolic pathways^5^. RaMP-DB 2.0, originally released in 2018, is a comprehensive relational database that integrates functional and other biologically relevant annotations for metabolites, genes, and proteins, where the latter are harmonized across the multiple sources HMDB^6–9^, Reactome^10,11^, WikiPathways^12,13^, and KEGG^14–16^ (through HMDB). Our intent for building RaMP-DB 2.0 was to provide up-to-date and comprehensive annotations that could be readily used to interpret metabolomic and multi-omic data by using the associated R package and web interface, or by integrating the publicly available relational database (e.g., available as a mySQL dump) directly into one’s own tools.

Among tools that harmonize multiple sources and support multi-omic analyses, RaMP-DB 2.0 is notable for its inclusion of multi-omic pathways from multiple sources, its ability to accept mixed ID types for genes, proteins, and metabolites, its focus on lipid and chemical structure annotations, and its ability to compute enrichment statistics using the aforementioned IDs and pathway (**Supplementary Table 1**). In fact, many tools rely only on a single pathway database, limiting the pathways that are considered in enrichment analysis. It is also worth noting that public knowledge sources and associated tools that draw information from multiple databases are oftentimes built for a specific purpose. For example, ConsensusPathDB^17^ focuses on physical interactions and supports interaction network exploration and single-omic enrichment analyses. IMPaLA^18^ focuses on multi-omic pathway analysis of transcripts/proteins and metabolites, but lacks special focus on metabolite functional enrichment in the form of lipid and chemical structure annotations. For each tool, only a portion of the original resources is parsed and included to meet the analysis needs. Analogously, RaMP-DB 2.0 was specifically built to support multi-omic biological pathway enrichment analysis and chemical class enrichment analysis of metabolites. RaMP-DB 2.0 is actively maintained, and its underlying processes are fully transparent with a GitHub page that allows users to ask questions, submit issues, and browse and download the source code for the R package, web interface, and database construction.

We present here on the recent enhancements to our RaMP-DB 2.0 platform. First and foremost, RaMP-DB 2.0 now utilizes a new semi-automated entity resolution method that verifies compound structural elements when mapping metabolite entries across different databases. This entity resolution method is augmented by manual curation to verify metabolite identifier mappings as well as implied relationships. The semi-automation process allows for frequent updates to be performed. Second, RaMP-DB 2.0 now also includes chemical structure and class annotations for metabolites. This information is useful for exploring the breadth of chemical classes and space covered by a collection of metabolites under study. Overrepresentation of chemical classes in a metabolite set of interest relative to a larger collection of quantified metabolites can also be calculated, providing complementary information to biological pathway enrichment, which typically informs on a much smaller fraction ofmetabolites^19^. Pathway and chemical enrichment analyses support the inclusion of a custom background (e.g. metabolites evaluated in a study). Lastly, RaMP-DB 2.0’s contents have been updated to reflect expansions of its constituent pathway databases. For example, the most recent ontologies from HMDB 5.0, including relevant portions of the new chemical functional ontology (CFO), are now included.

## MATERIAL AND METHODS

### Backend Code Base: Parsing and Harmonization

Python scripts acquire data from our primary data sources (**Table 1**) and parse annotations associated with pathways, common reactions, ontologies, chemical structures, and chemical classes. Each primary data source has a dedicated python class that fetches on-line data files to a local data store, then reads and parses the data source-specific input data and writes the key data to a collection of intermediate files of a standard format. A processing class reads the collection of all intermediate files for all data sources and populates a collection of entity classes that are organized into natural relationships between genes, metabolites, pathways, and related information. This data structure allows a well-controlled process of data harmonization, with error checking at redundancy checks. This data aggregation and harmonization step uses a configuration file that controls the database refresh and a python class for database loading to write a final set of files that are formatted to ease database upload.

**Table 1:**
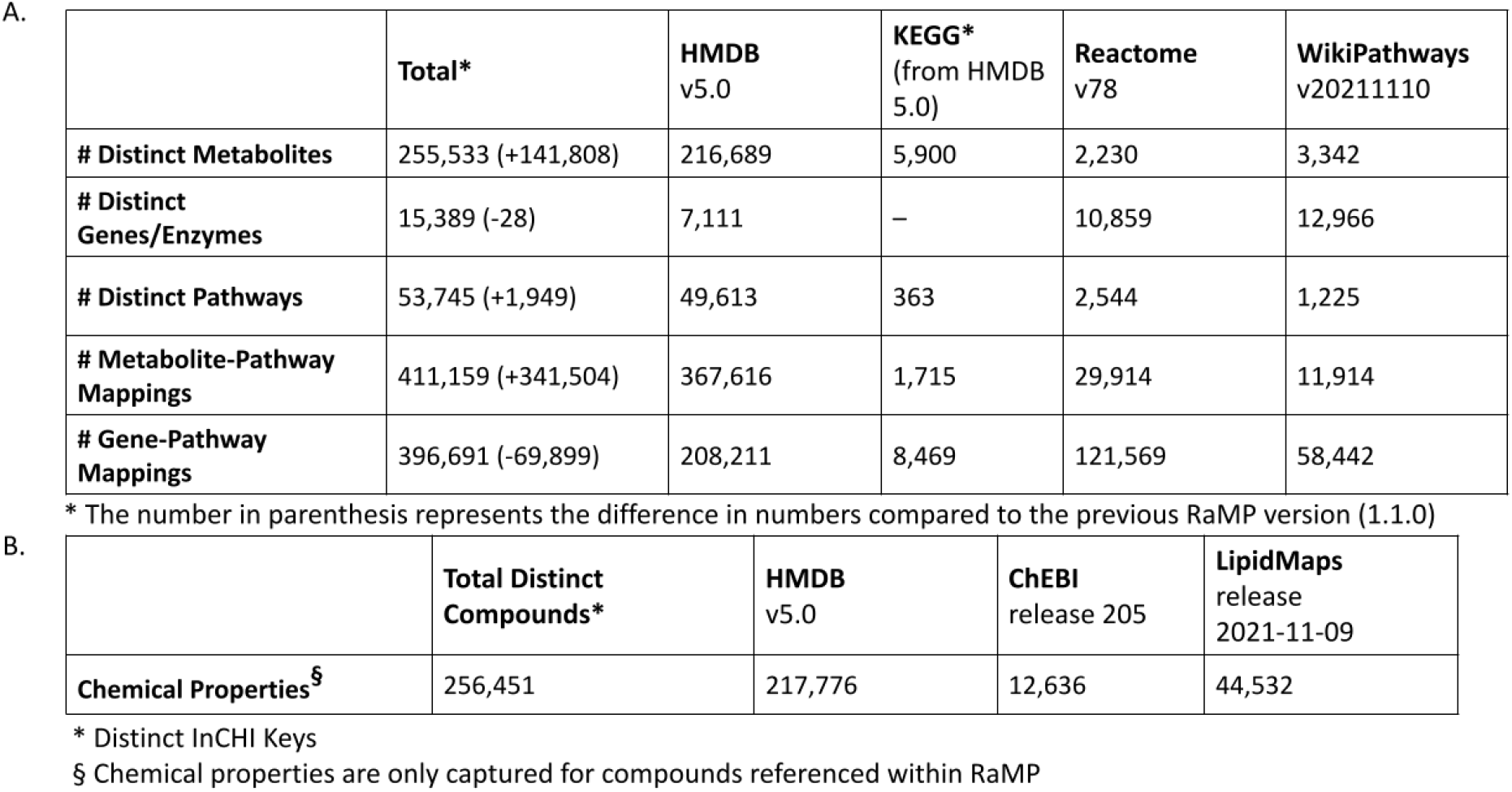
Number of analytes and pathways (A) and chemical properties (B) available through RaMP 2.0.

**Supplementary Figure 1C** diagrams the RaMP 2.0 database schema. The *analyte* table contains a list of internal and unique RaMP analyte IDs that correspond to genes, proteins, and metabolites. Meta-information associated with these analytes are featured in the *source* table and the *analytesynonym* table. The *source* table includes all IDs and common names that map to each RaMP analyte entity. *Analytesynonyms* provides 779,175 common name synonyms for the metabolites and genes in RaMP. Two mapping tables (*analytehaspathway* and *analytehasontology*) connect our analyte entity IDs with tables that contain information on pathways and metabolite annotations held in the *ontology* table. Separate tables contain metabolite information on chemical classes from HMDB (v5.0)^6–9^ and Lipid Maps^20,21^ (release 2021-11-09), and chemical properties from HMDB^6–9^ (v5.0), ChEBI^22–24^ (release 2021-11-03) and Lipid Maps^20,21^ (release 2021-11-09). Three tables contain meta-information that characterizes the current database build: the *db_version* table holds the build version and build timestamp of the entire RaMP 2.0 database, the *version_information* table holds information on each data source including the data sources version and release date, and the *entity_status_info* table contains a tally of current RaMP 2.0 entities within the build. Entities include counts for unique metabolites, genes/proteins, pathways, chemical property records for metabolites, and mappings between analytes and pathways in RaMP. The *catalyzed* table contains 1,542,009 associations between metabolites and genes that participate in metabolic reactions together.

All scripts to build the MySQL database are available through a public GitHub repository at https://github.com/ncats/RaMP-BackEnd. The resulting MySQL database dump is available for download through the public RaMP R package (https://github.com/ncats/RaMP-DB).

### Semi-Automated Entity Resolution

The RaMP-DB 2.0 database holds data from multiple data sources with each source contributing entities such as genes, proteins, and metabolites, and their annotations such as pathways, reactions, and ontologies. Associations between these entities and related annotations support interrogation of biology and chemistry, drawn across multiple sources, from any of these entity starting points. A depiction of the various entity types and their relationships is shown in **Supplementary Figure 1B**. This entity resolution allows users to input mixed ID types when performing batch queries, which is particularly relevant for metabolomic data that seldom report a single ID type. The different ID types supported for each analyte type are shown in **Table 2**.

**Table 2:** Supported ID types for analytes in RaMP

| Analyte type | Supported ID types |
| --- | --- |
| Metabolites | PubChem, HMDB, ChEBI, WikiData, ChemSpider, CAS, SwissLipids, LIPIDMAPS, KEGG, LipidBank, PlantFA |
| Genes/Proteins | HMDB, UniProt, Entrez, Gene Symbol, Ensembl, NCBIprotein, EN, WikiData, ChEBI |

To faithfully represent entities across various data sources, we have implemented a data model that accurately encapsulates associated entity meta-data (e.g. identifiers, synonyms, chemical properties, data source tags, etc.) and groups individual entities that represent the same molecule, as prescribed by the source data (**Figure 1**). Metabolite and gene entities are then connected to their corresponding biological pathway, chemical, and ontology annotations which contain names, external accessions, and links to other related gene/protein and metabolite entity records.

**Figure 1:**
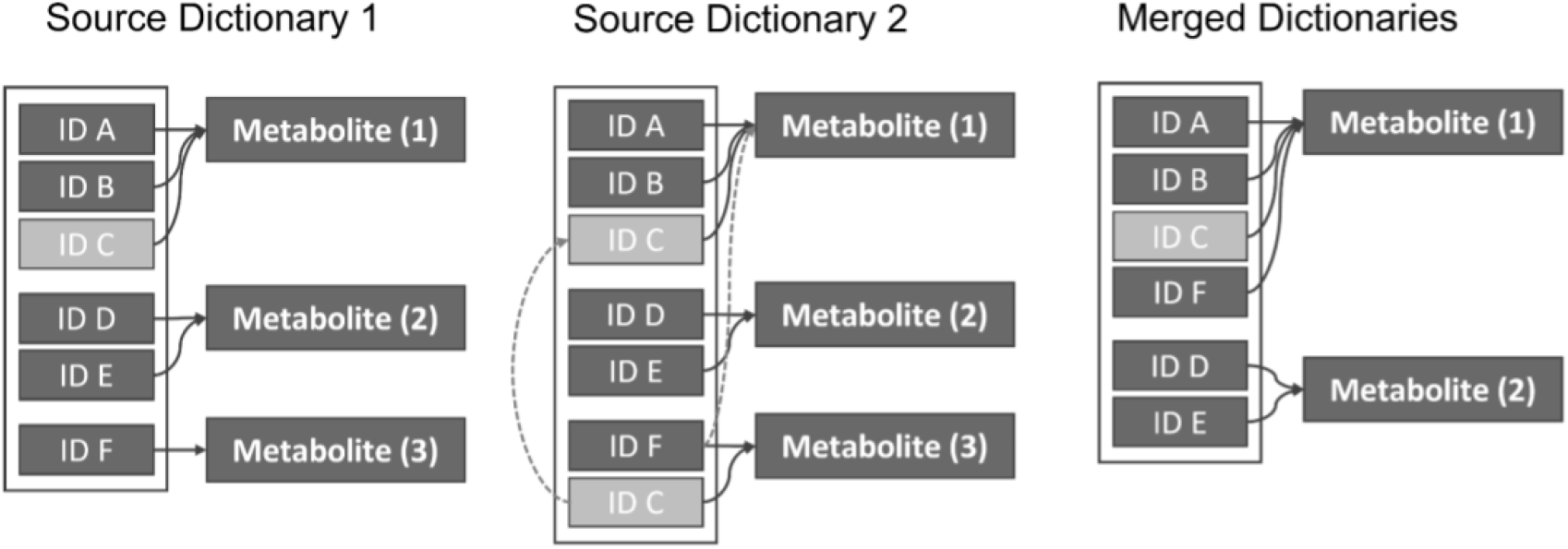
Approach to harmonization of names across different database sources. The figure depicts the mappings of IDs and metabolites from two different source databases (e.g. denoted as Source 1 and Source 2 Dictionary). Source 2 Dictionary has two instances of ID C, mapping to two different metabolites (1 and 3). In this case, Metabolites 1 and 3 will be merged and considered the same metabolite, which may or may not be accurate.

The data intake process starts with reading configuration files that instruct back-end processes to fetch data from external data sources. The source data files are then parsed with data-source specific parsers into intermediate consistently formatted files prior to combining and harmonizing data from the various data sources. The EntityBuilder class harmonizes the data by constructing all source data into a data model that deduplicates metabolites and holds all associations between metabolites and their chemical properties, genes, pathways and associated information. The harmonized data is written into final files prior to bulk loading into the relational database schema (**Supplementary Figure 1A)**.

Most data sources link metabolite entities to a collection of additional ID types, such as PubChem CID, ChemSpider, HMDB ID, ChEBI ID, and LIPID MAPS ID. ID cross-references of mappings help to suggest metabolites that are in common across different data sources. Two metabolites drawn from different sources that share a common metabolite ID are mapped to a common RaMP-DB 2.0 metabolite entity. Following the construction of these entities and relationships, the linked metabolites in the data model are verified by comparing molecular weights taken from the data source that publishes the identifier (HMDB, ChEBI, PubChem, Lipid Maps, or KEGG). Molecular weight variance of 10% or more from the lower molecular weight metabolite is flagged as a bad annotation and subsequently manually verified. During the automated assessment of all ID-based compound associations, suspect associations are flagged and exported to a list for manual curation. If it is deemed through subsequent manual curation that two IDs refer to different metabolites, then the metabolite ID pair goes into a list of associations to skip. After manual curation of all such issues, the data is built, skipping associations within the exclusion list. This results in both entities being represented, but skipping any merge suggested by the bad cross reference. This mis-mapping exclusion list is referenced on every subsequent database build. Reported discrepancies in the association between two metabolites IDs are added to the exclusion list as they are reported so that later database builds will use the latest curation patches.

### RaMP IDs

During the data harmonization and database loading, internal RaMP IDs are generated. These RaMP IDs are not intended to be used by the general user. Instead, they produce database-internal values for compound, gene, ontology, and pathway entities that act as keys that relate entities to one another, which are sourced from multiple sources, in the database. The values are used to reference RaMP entities within the database. While consistent within a database update, these IDs are not conserved across database versions. Entity IDs from external data sources are maintained as primary authoritative IDs to be used in result tables and in work derived from RaMP-DB 2.0 analyses. The RaMP-DB 2.0 primary analyte source information table and pathway table maintain a field that tracks the associated primary data source so that all RaMP-DB 2.0 entities maintain a record of data provenance.

### R Package

RaMP-DB 2.0 functions are annotated with roxygen v7.1.2 blocks to generate Rd help files, with working examples that can be opened in R. The package includes an extensive Vignette tutorial (https://ncats.github.io/RaMP-DB/RaMP_Vignette.html) that features the primary functions available within the package. Instructions on the GitHub page describe setting up the MySQL database locally and installing the RaMP-DB 2.0 R package. Importantly, the RaMP-DB 2.0 MySQL full database dump is included in the RaMP-DB 2.0 Package GitHub site and can also be explored independently of the R package. Like all parts of RaMP-DB 2.0, the RaMP-DB 2.0 Package is open source R, housed at https://github.com/ncats/RaMP-DB.

### Pathway and Chemical Enrichment Analysis

We have implemented functions for testing enrichment of pathways and chemical classes in a list of metabolites and/or genes and proteins. Enrichments are calculated using a Fisher’s Exact test, based on a 2×2 contingency table. These contingency tables are used to test the null hypothesis that the number of altered metabolites belonging to that pathway is less than or equal to the number expected by random chance. Where *n* is the total number of metabolites in the reference background, *a* is the count of differentially expressed analytes in the pathway being tested, *b* is the count of background analytes in pathway being tested *c* is the count of differentially expressed analytes outside of the pathway being tested, and *d* is the count of background analytes outside the pathway being tested, the Fisher’s *p-*value is calculated using **Equation 1**.

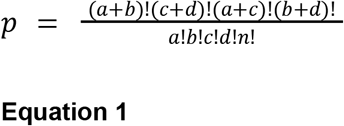

The reference background for the test can be defined in several ways. First, it can be defined as the number of metabolites contained in the pathway database that the pathway being evaluated is found in (e.g in this case, the entire RaMP-DB 2.0). Second, the reference background can be described as the full set of metabolites that were identified in the study. Last, it can be defined as the collection of metabolites that are known to be present in the biospecimen of interest (e.g. urine or blood metabolites only).

As in RaMP-DB 1.0, the RaMP-DB 2.0 R package includes a function for clustering pathways based on the overlap of their constituent analytes. This function is useful for easing interpretation of pathway analysis results, as RaMP-DB 2.0 contains many pathways that significantly overlap with other pathways contained in the database. As a new feature to RaMP-DB 2.0, we have implemented a “lollipop plot” function for plotting pathway analysis results that displays pathway cluster membership, database of origin for pathways, the number of altered analytes mapping to that pathway, and Fisher’s p value with multiple test correction of choice applied.

## RESULTS

### RaMP-DB 2.0 ecosystem

RaMP-DB 2.0 has two main components: 1) building of the back-end MySQL database that draws from multiple sources, and 2) an R package that supports queries and analyses using RaMP-DB 2.0. The RaMP R package supports four different batch query types along with two different enrichment analyses (**Figure 2A)**. Queries are available for lists of genes, proteins, and metabolites, returning pathways, biochemical reactions, and/or ontologies that contain those analytes. Users may also query a list of pathways and return analytes associated with those pathways. Lastly, for lists of metabolites only, users may query ontologies from HMDB, chemical structure, or chemical class information. Mappings returned from pathway and chemical class queries can be leveraged for functional enrichment analysis using the Fisher’s exact test.

**Figure 2:**
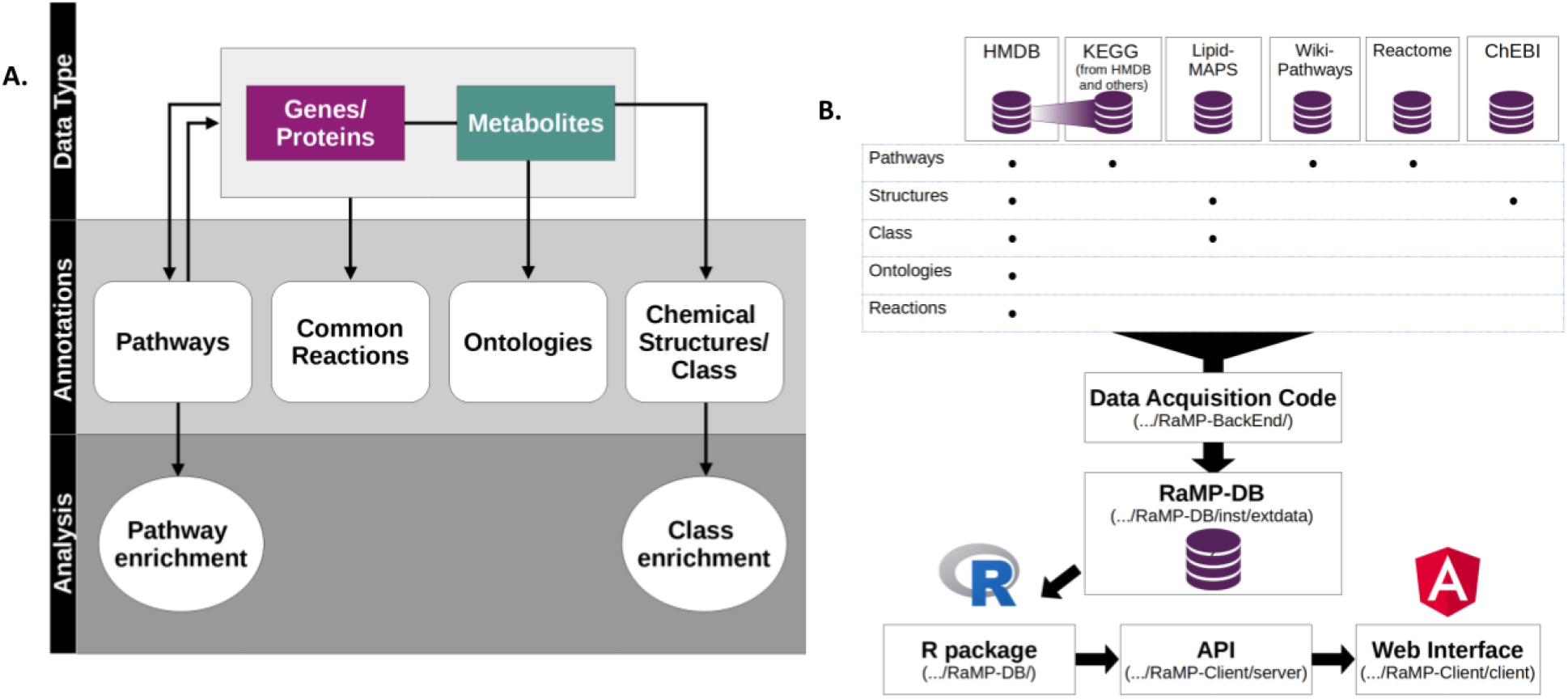
Overview of RaMP-DB, including its supported queries (A) and content (B).

Queries supported include: 1) Retrieve Analytes From Input Pathway(s); 2) Retrieve Pathways From Input Analyte(s); 3) Retrieve Metabolites from Metabolite Ontologies; 4) Retrieve Ontologies from Input Metabolites; 5) Retrieve Analytes Involved in the Same Reaction; 6) Retrieve Chemical Properties from Input Metabolites; 7) Retrieve Chemical Classes from Input Metabolites. The package also supports pathway and chemical class enrichment analysis of metabolites/genes/proteins and metabolites, respectively.

### RaMP-DB 2.0 Structure and Contents

The data for RaMP-DB 2.0 is drawn from six distinct sources (**Figure 2B**), and parsed using python scripts available at https://github/ncats/RaMP-BackEnd. The parsed data are organized into RaMP-DB 2.0, an analyte-centric relational database containing 13 tables (**Supplementary Figure 1C**). As in previous iterations, the relational structure offered by MySQL is designed for efficient retrieval of annotations related to a list of analytes of interest input by the user.

RaMP-DB 2.0 incorporates pathway annotations from four popular public metabolite pathway databases: HMDB^6–9^, Reactome^10,11^, WikiPathways^12,13^, and KEGG^14–16^ (through HMDB) (**Table 2**). RaMP-DB 2.0 contains pathway associations for both genes/proteins and metabolites. In total, following entity resolution, RaMP-DB 2.0 contains 151,526 metabolites, 14,362 genes, and 52,573 pathways (see **Semi-Automated Entity Resolution in Methods** for details on entity resolution).

New additions to RaMP-DB 2.0 include ontologies derived from HMDB (v5.0)^6–9^, chemical structure and class information. Most notably, the inclusion of 43,448 lipids from LIPID MAPS has greatly expanded the number of lipids for which information is available in RaMP-DB 2.0. RaMP-DB 2.0 now contains 170,654 chemical structures associated with its collection of metabolites. These structures, obtained from HMDB (v5.0)^6–9^, ChEBI (release 2021-11-03)^22–24^ and LIPID MAPS (release 2021-11-09)^20,21^, provide a rich source of information for chemical compounds of interest and can be used as a basis for cheminformatics analysis. RaMP-DB 2.0 also contains chemical class, superclass, and subclass as dictated by the ClassyFire^25^ taxonomy and the LIPID MAPS database, which can also be used as a basis for chemical class enrichment analysis. Lastly, RaMP-DB 2.0 contains a collection of 807,362 metabolic enzyme/metabolite reactions and 699 ontologies from HMDB^6–9^ (v 5.0).

Integrating data from multiple resources expands the number and type of annotations for analytes. With RaMP-DB 2.0, an analysis of overlapping content in each of the constituent databases is thus possible, highlighting the value of a united database for these annotations. Out of the 151,526 unique metabolites found in RaMP-DB 2.0, 103,616 metabolites (68%) are only found in one of the source databases (99% of these unique metabolites are derived from HMDB, **Figure 3**). Out of 56,840 metabolites that have at least one pathway association in RaMP, 12,758 are only found with pathway mappings in one source database (22%). Similarly, out of 14,362 genes/proteins in RaMP-DB 2.0, 3,323 genes/proteins (23%) are unique to their source database.

**Figure 3:**
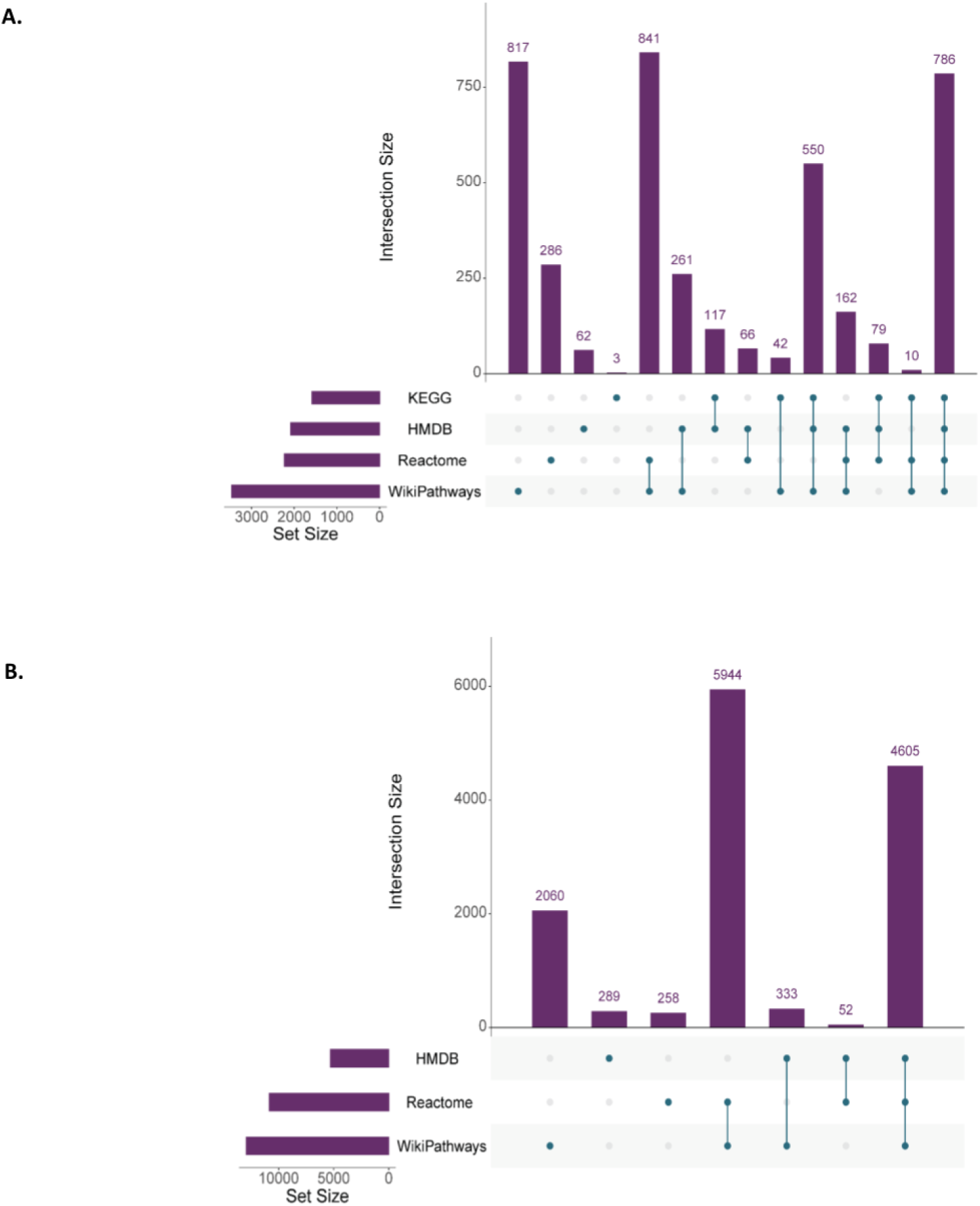
Overlap in content among source databases. A) Overlap in metabolites associated with pathways provided by each source database in RaMP. B) Overlap of genes associated with pathways provided by databases in RaMP. The filled circle(s) underneath each bar in the plots demonstrate the source databases that the analyte counts are drawn from.

### Resolving Entity Mismappings

Because RaMP-DB 2.0 relies on accurate ID mappings for a metabolite in one resource (e.g. metabolite X mapping to multiple IDs YYY), mis-mappings could create errors in linking one metabolite to another from different sources. As an example, we found an occurrence where a data source provides a metabolite record for a diglyceride and provides corresponding valid PubChem ID (**Supplementary Figure 2**). Another metabolite record from the same source represents 11-Oxo-androsterone glucuronide but has an external ID reference to the PubChem diglyceride record. Naively following ID associations would collapse these two metabolites into one entity due to the common PubChem CID. In this case our steroid-based compound has a molecular weight of 480.55 Da, while the diglyceride has a molecular weight of 681.12 Da so the error in linking was identified automatically through molecular weight comparison.

Notably, these mis-mappings would propagate to errors in pathways or other annotation mappings and introduce false positives in enrichment analyses. To address this issue, we have developed a heuristic based on MW weight to automatically flag potential mis-mappings that could occur between database sources (described in **Semi-automated entity resolution** above). These flagged mis-mappings are then manually investigated and fixed as appropriate. Mis-mappings are recorded and used to automatically correct future updates of RaMP-DB 2.0.

RaMP-DB 2.0 curation revealed a total of 955 distinct metabolites involved in incorrect associations linking disparate molecules. Within the 955 metabolites, 351 metabolites had incorrect ID-based cross reference associations to 604 distinct metabolites. Curation eliminated the propagation of these errors and the resulting mis-associations that could arise between these metabolites and pathways.

## DISCUSSION

To the best of our knowledge, RaMP-DB 2.0 is the only knowledge source and associated tool that supports batch queries of analyte annotations, multi-omic pathway and chemical class enrichment analysis with ability to input mixed ID types, and batch queries of pathway and chemical annotations using mixed identifier schemes for both genes and metabolites. Incorporating multiple sources into enrichment analyses greatly expands the mappability of analytes to pathways, thereby enhancing the user’s ability to functionally interpret complex data. RaMP-DB 2.0 verifies the accuracy of mapping analyte entities across its various source databases using a semi-automated process followed by manual curation. The updated RaMP-DB 2.0 now includes 151,526 metabolites, 13,927 genes, 52,573 pathways, 408,232 mappings between metabolites and pathways, and 463,447 mappings between genes and pathways. Improving the accuracy of mappings between analyte identifiers will increase the accuracy of downstream insights gleaned from data.

New features of the updated version of RaMP-DB 2.0 include the ability RaMP-DB 2.0of users to perform chemical class enrichment analysis on metabolites of interest. Other recent efforts in metabolomic software development have noted that chemical class and substructure enrichment analysis can provide functional insight where pathway annotations are unavailable, as chemical structure annotations for metabolites typically offer better coverage^19,26^. The primary benefit of class enrichment is the superior coverage of chemical class annotations available for metabolites, thanks to the ClassyFire taxonomy^25^. In RaMP-DB 2.0, each metabolite has a superclass, class, and subclass designation, while only 36.6% of metabolites have at least one pathway annotation associated with them. Better coverage allows for the incorporation of more experimental information into test results. Class enrichment also allows for the testing of different hypotheses than pathway analysis. For example, in studies where the objective is to identify putative therapeutic targets, discovery of altered classes can suggest enzymes acting upon generic species of that class as potential inhibition targets^27^. Integrating this functionality into RaMP-DB 2.0 gives users another option for gaining functional insight into their data.

We also note the importance of using an appropriate background/reference list of analytes for pathway enrichment analysis^28,29^. Typically, “background” metabolites used for the Fisher’s exact test are defined as all metabolites in the original database the pathway being tested was derived from. However, an alternative definition is the list of all metabolites identified in a study. Many Fisher’s pathway analysis tools such as DAVID^30^ and MetaboAnalyst^31,32^ enable users to select their choice of background. Accordingly, we have implemented the option for either background selection in RaMP 2.0. We have also implemented a novel third option for backgrounds, comprising all metabolites known to occur in a given biospecimen (as determined by HMDB ontology). Example biospecimen types include “Adipose Tissue”, “Blood” and “Heart”. We note that using a broad background could include metabolites that should be excluded from the analyses because they are absent in the biospecimen under study, or were not detected for some reason (e.g. failure to ionize or exclusion due to the extraction protocol used). As such, a more appropriate hypothesis to test is to use a custom background of only those metabolites detected in the study, or those appropriate for the biospecimen of interest. A recent study of metabolomics pathway analysis strategies confirms that choice of background in the Fisher’s exact test exerts large effects on the list of significant pathways returned by the Fisher’s exact test^33^. Enabling users to choose their background based on the information available could thus lead to more reliable outputs for enrichment analysis. Nonetheless, we preserve the option of using a database background, as the list of all metabolites identified in an experiment is not always readily available to the researcher.

Despite our recent enhancements, RaMP-DB 2.0 has some limitations, particularly regarding its coverage and harmonization. Currently, RaMP-DB 2.0 is limited to human pathways, although metabolite coverage does include microbial, food, and other exogenous metabolites. Furthermore, the harmonization of metabolites across the different resources is not fool-proof. Metabolite identification in large-scale metabolomic experiments is still an unresolved issue, and experiments often yield a mix of different levels of certainty and structural resolution in identified metabolites^34–39^. These factors are not taken into account during harmonization or mapping of metabolites across sources. Users should thus carefully assess their input list of metabolites, particularly for enrichment analysis and double check that mapping of metabolites is correct. This process is facilitated by the modular design of running enrichment analyses, as described above, which requires users to review database mappings from their input IDs as well as pathway mappings before seeing enrichment results. In all cases, we recommend that users input IDs, rather than names for analyses (this is the default implementation by design).

As a future direction, RaMP-DB will contain reaction-level pathway network information drawn from KEGG and Reactome to allow for the use of topological pathway analysis algorithms on metabolomic data. A notable drawback of Fisher’s exact test is that it treats all metabolites in a pathway as equivalent parts of a set^40^. This is a misrepresentation, as pathways are a collection of metabolites undergoing reactions that result in some signal or biochemical product. More specifically, a metabolite is less likely to exhibit correlated abundance with another random metabolite in the same pathway as it is with a known reaction partner in the same pathway. While topological pathway analysis methods are more mature in transcriptomic applications^41^, implementations exist in the metabolomic field as well. For example, in MetaboAnalyst^32^, the pathway topological analysis module uses relative betweenness centrality and out degree centrality network metrics to assign relative importance scores to metabolites in a pathway using a network representation wherein metabolites are nodes and reactions are edges. Intuitively, this score is used to assign greater weight to metabolites that are more central in a pathway, meaning they are well-connected to other parts of the pathway and more likely to exert influence downstream. Ultimately this results in more “important” metabolites contributing more to pathway perturbation scores than less central metabolites. We also anticipate the inclusion of more data types for genes and proteins such as gene ontology annotations or additional reactions from Rhea^42^. As with other annotations in RaMP-DB 2.0, functions for batch querying and enrichment analysis of new annotations will be implemented. Lastly, the R package relies on a local instance of RaMP rather than calling upon API, which forces users to install and store their own copy of the database. In the future we intend to remove this limitation.

## CONCLUSIONS

RaMP-DB 2.0 is a multi-sourced relational database comprising pathway and chemical annotations for metabolites, genes and proteins. An improved resolution of metabolite and gene/protein mappings across the databases has been implemented and is supplemented with manual curation. Associated and improved R package and web user-friendly interface have been constructed to query the database and perform chemical and pathways enrichment analyses. All steps of RaMP-DB 2.0 are reproducible with all the code used to build or use the database publicly available in GitHub.

## DATA AVAILABILITY

RaMP-DB 2.0 is an open source project available in the GitHub repository (https://github.com/ncats/ramp-db), which includes code for the R package and code for constructing the RaMP-DB 2.0 mySQL database, as well as the latest RaMP-DB 2.0 dump.

Python code for building the RaMP-DB 2.0 database can be found at https://github.com/ncats/RaMP-Backend.

## ACKNOWLEDGEMENTS

Thank you to the beta testers for providing early feedback on the development of RaMP-DB 2.0

## SUPPLEMENTARY TABLES AND FIGURES

**Supplementary Table 1:** Publicly available tools for functional enrichment analysis of metabolomic and transcriptomic data

| Tool | Reference Database | Multi-omic pathway analysis with mixed ID input | Multi-omic pathway analysis using multiple knowledge sources | Inclusion of extensive chemical/lipid annotations |
| --- | --- | --- | --- | --- |
| **RaMP** | HMDB, Reactome, Wikipathways | Yes | Yes | Yes |
| **MetaboAnalyst** | KEGG, Pathbank | No | Yes | No |
| **3Omics** | KEGG, HumanCyc | No | Yes | No |
| **IMPaLA** | 11 total including KEGG, Reactome, BioCyc, Wiki-Pathways, SMPDB | No | Yes | No |
| **ConsensusPathDB** | 30 different sources including Reactome, KEGG, WikiPathways and HumanCyc | No | Yes | No |
| **MetaBox** | KEGG | No | No | Yes |
| **PathNet** | KEGG | No | No | No |
| **GO-Elite** | GO, WikiPathways, PathwayCommons, KEGG | Yes | Yes | No |

**Supplementary Figure 1:**
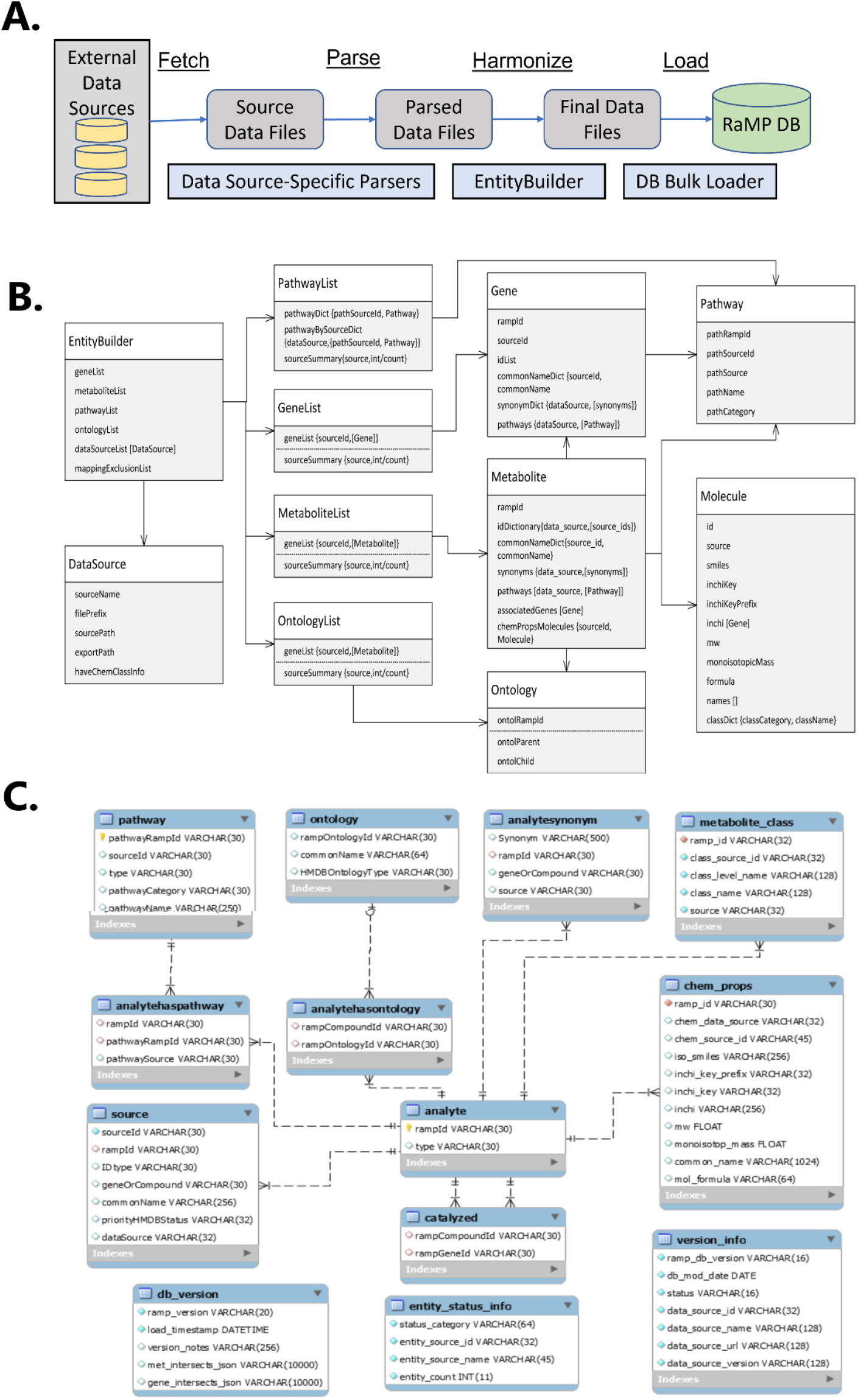
**A)** Overview of the process underlying the backend code used to create RaMP **B)** Data types stored within RaMP-DB 2.0 and their relationships **C)** RaMP-DB 2.0 Schema (and data models)

**Supplementary Figure 2:**
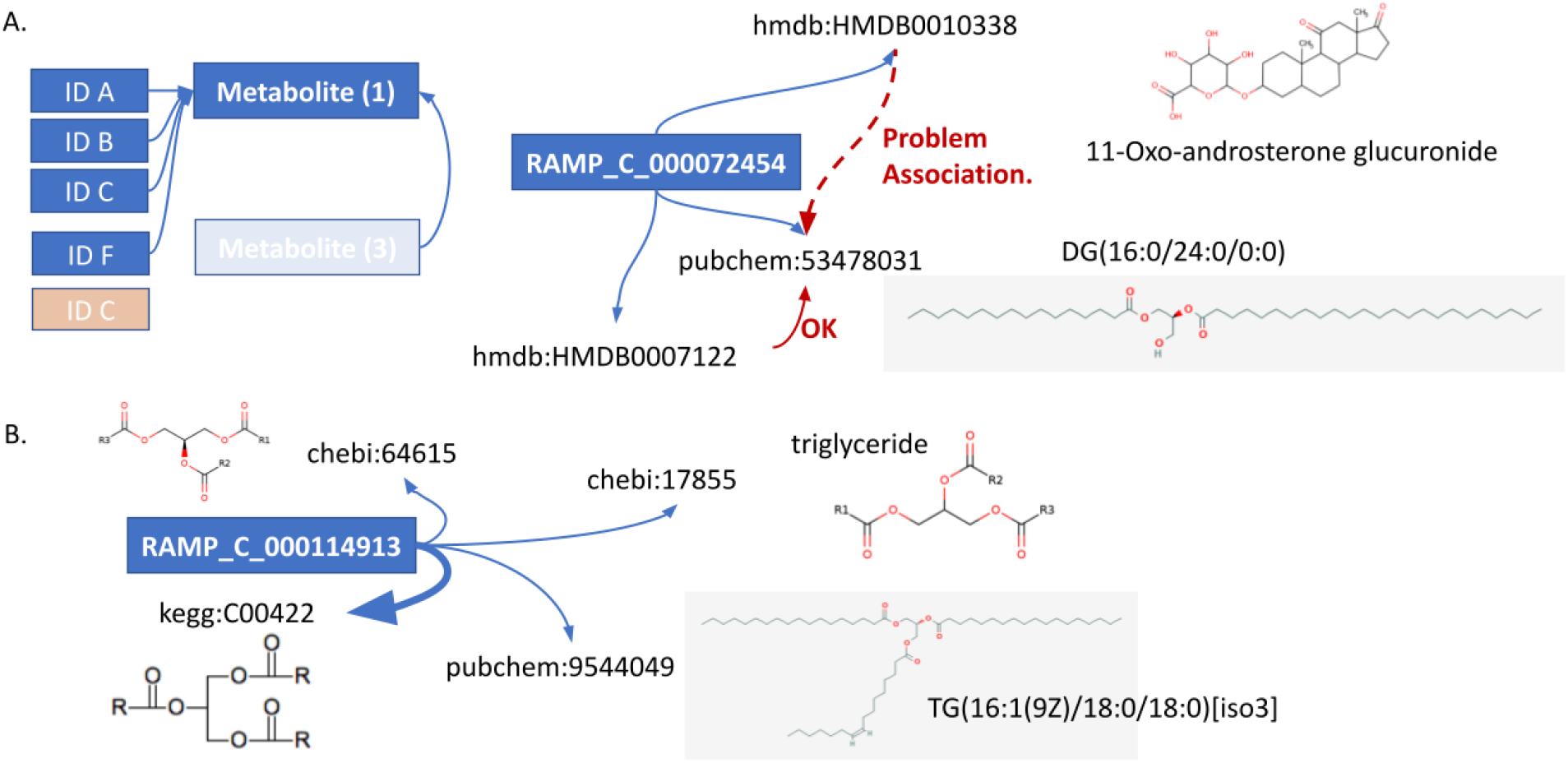
Example errors in mappings. A) Two completely different structures are mapping to the same PubChem ID. B) Specific structures are mapping to generic structures (triglycerides with R groups rather than specific chains). 118 of 125 HMDB triglycerides map to the generic KEGG triglyceride ID (C00422).

